# PLASTICITY OF RESPONSE PROPERTIES OF MOUSE VISUAL CORTEX NEURONS INDUCED BY OPTOGENETIC TETANIZATION *IN VIVO*

**DOI:** 10.1101/2023.10.01.556796

**Authors:** A.A. Osipova, I.V. Smirnov, M.P. Smirnova, A.A. Borodinova, M.A. Volgushev, A.Y. Malyshev

## Abstract

Heterosynaptic plasticity, along with Hebbian homosynaptic plasticity, is an important mechanism ensuring stable operation of learning neuronal networks. However, whether heterosynaptic plasticity occurs in the whole brain in vivo, and what role(s) in brain function in vivo it could play, remains unclear. Here, we used an optogenetics approach to apply a model of intracellular tetanization, which was established and employed to study heterosynaptic plasticity in brain slices, to study plasticity of response properties of neurons in mouse visual cortex in vivo. We show that optogenetically evoked high-frequency bursts of action potentials (optogenetic tetanization) in principal neurons of the visual cortex induce long-term changes of responses to visual stimuli. Optogenetic tetanization had distinct effects on responses to different stimuli: responses to optimal and orthogonal orientations decreased, response to null direction did not change, and responses to oblique orientations increased. As a result, direction selectivity of the neurons decreased, and orientation tuning became broader. Since optogenetic tetanization was a purely postsynaptic protocol, applied in the absence of sensory stimulation, and thus without association of presynaptic activity with bursts of action potentials, the observed changes were mediated by mechanisms of heterosynaptic plasticity. We conclude that heterosynaptic plasticity can be induced in vivo and propose that it may play important homeostatic roles in operation of neural networks by helping to prevent runaway dynamics of responses to visual stimuli and to keep the tuning of neuronal responses within the range optimized for encoding of multiple features in population activity.

## INTRODUCTION

Synaptic plasticity represents the cellular basis of learning. Synaptic plasticity is not a uniform phenomenon, rather, multiple forms and mechanisms have been described and studied [1,2]. Synaptic plasticity can be segregated into homosynaptic, which requires activation of the presynapse for the induction, and heterosynaptic, which does not require activity at the presynapse for the induction [3,4]. In this paper we are following this use of the terms homosynaptic and heterosynaptic, which is conventional for research of plasticity in mammalian nervous system [5]. Relation to the terminology used in invertebrate research [6–8] will be considered in the discussion.

Distinct forms of plasticity play diverse functional and computational roles. Homosynaptic plasticity can be Hebbian and associative, and thus mediate associative learning. Its mechanisms are studied in most detail, in diverse experimental preparations, including plasticity of response properties and receptive fields of neurons in visual cortex *in vivo* [9–11]. Heterosynaptic plasticity is a necessary component of learning networks, helping to robustly prevent runaway dynamics, enhance synaptic competition, and increase the contrast of synaptic weight changes [4,12–14]. It received less attention than homosynaptic plasticity, and was studied predominantly in slice preparations, which allow for better control of stimulation and especially for the absence of presynaptic activity during the induction. In slices from the hippocampus, heterosynaptic long-term depression (LTD) accompanying tetanus-induced long term potentiation (LTD) but occurring at non-activated synapses was first described by Lynch and colleagues [5]. A feasible trigger for this form of heterosynaptic plasticity are bursts of action potentials [15,16] which are generated during the induction of homosynaptic LTP and back-propagate into the dendritic tree [15]. On a local scale, heterosynaptic plasticity could be induced by the diffusion of diverse signaling molecules within the dendrite or in the extracellular space, causing a Mexican hat-shaped profile of LTP and LTD around the location of the tetanized synapses [17–19], breakdown of input specificity of plastic changes [20] and even a spread of plasticity to closely located but non-stimulated neurons after pairing [21]. An established experimental paradigm to study heterosynaptic plasticity is intracellular tetanization - bursts of spikes induced by depolarization pulses without presynaptic stimulation. Plastic changes induced by such purely postsynaptic protocol can be interpreted as heterosynaptic. Intracellular tetanization paradigm has been applied to study plasticity in diverse cells and preparations: pyramidal neurons in the hippocampus [16], granular neurons of the dentate gyrus [22], excitatory and inhibitory neurons of the neocortex [23–28], and even identified neurons of the common snail [29,30]. In contrast to in vitro preparations, evidence for heterosynaptic plasticity *in vivo* is sparse, in part because of difficulties to control for the absence of presynaptic stimulation during plasticity induction. In the visual cortex, heterosynaptic changes were described in experiments using two-photon microscopy imaging of calcium signals in spines. In mouse visual cortex, pairing the activation of spines using visual stimulation with postsynaptic spikes induced LTP, and in other spines at the same neuron heterosynaptic LTD could be induced [31]. While absence of presynaptic activation during the induction was not controlled for and cannot be excluded, especially considering huge size of receptive fields studied and strong visual stimulation used, these data indicate that heterosynaptic plasticity can be induced in mouse visual cortex *in vivo*.

Here we set to resolve the question of whether heterosynaptic plasticity can be induced *in vivo*. To circumvent the problem of presynaptic activation of the test inputs during the induction we used a purely postsynaptic induction protocol of intracellular tetanization – bursts of postsynaptic spikes evoked using optogenetic stimulation of the recorded neuron without presynaptic activation. To achieve lasting recordings and to avoid another potential problem of ‘plasticity wash-out’ we employed juxtracellular recording. We show that, in the principal neurons of the mouse visual cortex, bursts of high-frequency action potentials without presynaptic activation can induce long-term changes of responses to visual stimuli and a change of directional preference of neurons. Because there was no visual stimulation and presynaptic activation during the induction, the observed changes of responses were mediated by heterosynaptic plasticity. We conclude that heterosynaptic plasticity can be induced *in vivo* and propose that it could play a vital role in maintaining stability of operation and functionality of learning networks *in vivo*.

## METHODS

### Animals

Experiments were performed on adult C57Black/6 mice, 2–4 months of age (Pushchino Breeding Center, Branch of the Shemyakin–Ovchinnikov Institute of bioorganic chemistry of RAS). All experimental procedures were in accordance with the 2010/63/EU directive for laboratory animals. The research protocol was approved by the Ethical Committee of the Institute of Higher Nervous Activity and Neurophysiology, Russian Academy of Sciences.

### AAV virus production and injection

Virus injection was made under isoflurane anesthesia. Dexamethasone (2 mg/kg, intracutaneous) was used to prevent inflammation after the surgery. AAV2-CaMKIIa–oChIEF-EGFP virus was produced at local facilities and purified on HiTrap heparin column (Cytiva). Virus was slowly injected intracranially in the right hemisphere at stereotaxic coordinates (A/P = -2.8, M/L = 1.4 from Bregma), 200 μm below the brain surface (1 μl in concentration of 1.49E+12 vg/ml in PBS, injection rate 0.06 μl per minute).

### Surgery

Recording experiments were made 2-4 weeks after viral injection. During surgery and recording, the animal was kept under urethane anesthesia (0.5-1.5 g/kg). Additionally, atropine (5 mg/kg) and dexamethasone (2 mg/kg) were administered subcutaneously to reduce secretion and edema. During the surgery, the head of the animal was fixed in the head holder SMG-4 (Narishige, Japan). To improve the signal-to-noise ratio during intrinsic imaging, the scull over V1 was thinned out. The head plate was affixed to the skull with dental cement on cyanoacrylate glue. After one hour of recovery from surgery, the mouse was placed in the experimental setup.

### Visual stimulation

Visual stimuli were presented on a LED monitor (mean luminance 45 Cd, 60 Hz) placed 25 cm from the mouse, covering ∼70° of visual space. Visual stimuli, drifting sine-wave gratings (0.04 cpd, 2 Hz, 12 directions) were generated using PsychoPy [32]. During electrophysiological recording, drifting gratings of 12 orientations were presented in pseudorandom order, each grating presented for 2 seconds with 0.5 second interval (gray screen) between the stimuli. Sets of stimuli moving in all 12 directions were repeatedly presented during the recording.

### Intrinsic optical imaging

Prior to electrophysiological recording, the retinotopic projection of the monitor onto V1 was determined using intrinsic optical signal (IOS). IOS was recorded using a CMOS camera (BFS-U3-17S7M-C, FLIR, USA; 40 fps) and Micro-Management software. During the first 2 seconds of the trial, background frames were collected while a grey screen was presented, then visual stimuli (drifting gratings, 0.5 second for each of 12 orientations) were presented for 6 seconds, and then grey screen was presented for 12 seconds. A total of 30 of such 20 second trials were collected. Each frame was then normalized to mean intensity and the average background frame was subtracted from the averaged response frame. Using the resulting image, the center of the retinotopic projection of the screen was determined as the area with the lowest intensity. A small craniotomy (300-500 μm) was then made in the center of the area corresponding to the retinotopic projection of the screen.

### Electrophysiological recording

For electrophysiological recordings we used juxtracellular technique. Borosilicate glass electrodes were filled with Hanks’ Balanced Salt Solution (HBSS, 138 mM NaCl, 1.26 mM CaCl_2_, 0.5 mM MgCl_2_, 0.4 mM MgSO_4_, 5.3 mM KCl, 0.44 mM KH_2_PO_4_, 4.16 mM NaHCO_3_, 0.34 mM Na_2_HPO4, 10 mM glucose, and 10 mM HEPES at pH 7.4) and had a resistance of 4–12 MΩ. An optic fiber (200 μm core) was inserted in the recording electrode using a specialized electrode holder (Optopatcher, A-M Systems, USA) and connected to a 470 nm LED (ThorLabs, USA).

Light intensity, measured at the tip of the optical fiber with the maximal LED power, was around 5 mW. During the cell search short light pulses were constantly delivered through the optic fiber. Only neurons that generated spikes in response to every optogenetic stimulus (i.e. oChIEF expressing neurons) were taken in the experiment. The experiment was performed as follows. First, control responses to moving gratings were recorded for 15–40 minutes. Then visual stimulation was stopped, and recorded neuron was either optogenetically tetanized (tetanus group) or was recorded without any stimulation for 5-6 min (control group). Control experiments were performed on the non-transfected mice. Optogenetic tetanization consisted of 5 series of 10 trains of light pulses delivered through the optic fiber. Inter-series interval was 1 min and inter-train interval was 1 s. Every train consisted of 8-10 pulses with the frequency of 50-100 Hz and pulse duration of 8 ms. The frequency and the number of pulses were adjusted individually for each cell in order to induce APs with maximal possible frequency (70 – 100 Hz). During tetanization no visual stimuli were presented. In the control group, there was a pause in the visual stimulation for 5-6 min, corresponding to the duration of optogenetic tetanization. After the tetanization (or a pause in stimulation in the control group), visual stimulation was resumed and responses to moving gratings were recorded for at least 1 hour. Recordings were made using a Multiclamp 700B amplifier (Molecular Devices, USA) with PClamp 10 software (Molecular Devices, USA), the signals were filtered at 300-10,000 Hz and digitized at 20 kHz (Digidata 1550 Series, Molecular Devices, USA).

### Labeling

To label recorded neurons, 1% neurobiotin was added to the electrode solution in some experiments (Vector Laboratories, USA). To deliver neurobiotin into the neurons 500 ms positive current pulses at 2 Hz frequency were applied during the last 10 minutes of the recording. The amplitude of the current pulses was 2-10 nA, adjusted individually for each neuron. After allowing neurobiotin to diffuse in the cell for one hour, mice were perfused with 10% PFA, then brain was extracted and post-fixed in 4% PFA overnight. 50-mkm slices were made using a vibratome (Leica 1100, Germany). Neurobiotin was visualized by incubating slices in 1:250 Streptavidin-Rhodamin (ThermoFisher, USA) solution in PBS. The slices were examined using laser confocal Cerna-based microscope (Thorlabs, USA).

### Data analysis

Data analysis was done using custom scripts in Python3. APs were detected by a threshold which was set individually for each cell. To calculate the F1/F0 ratio, the peristimulus time histogram of visual responses was fitted with a 2 Hz sine wave (corresponding to the spatial frequency of the moving gratings) and then the amplitude of the fitted sine wave was divided by the average amplitude of the response. The direction and orientation selectivity indices (DSI and OSI) were calculated as follows:

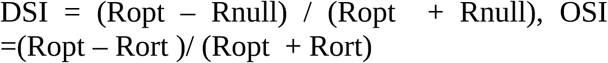

where, Ropt – response to the visual stimulus of the optimal orientation and direction, Rnull – response to the visual stimulus of optimal orientation but moving in the opposite to the optimal direction, and Rort – averaged response to the visual stimuli of orthogonal to optimal orientation, moving in both directions. Values are presented as mean + SD.

## RESULTS

The aim of our work was to investigate whether a purely postsynaptic challenge – bursts of action potentials without presynaptic stimulation (unpaired tetanization), can induce changes of response properties of neurons in the mouse visual cortex *in vivo*. To achieve stable long-term recording from individual neurons without compromising the integrity of their membrane while having a possibility to stimulate them through the recording pipette, we chose the technique of juxtacellular recording. However, a pilot series of experiments showed that passing through a juxtacellular electrode current pulses of high amplitude, necessary for the induction of high-frequency bursts of action potentials, resulted in membrane rupture. To circumvent this problem, we decided to use an optogenetic approach for tetanization of neurons. Classical channelrhodopsin2, due to the pronounced sensitization of responses during rhythmic stimulation, allows to induce controlled bursts of action potentials at frequencies up to 20–30 Hz, but not higher [33]. Therefore, we used the fast channelrhodopsin oChIEF, which allows efficient optogenetic stimulation of neurons at frequencies around 100 Hz [34], thus letting us to reproduce conventional plasticity-induction protocols. To transfect neurons in the mouse primary visual cortex with fast channelrhodopsin oChIEF we injected the AAV2-CamKII-oChIEF-Venus virus medio-caudal to the intended recording site. Recordings were started at least 2 weeks after the injection. Morphological control made after electrophysiological experiments revealed strong fluorescence in all cortical layers around the site of injection (Fig 1A). At the recording sites, 1-2 mm away in lateral-posterior direction from the injection site, fluorescence pattern was not uniform across the layers. There was a pronounced fluorescence in layers 1, 2/3 and 5a of the visual cortex and in the white matter (Fig. 1A). In layer 4 fluorescence was very weak, only in the dendrites of the layer 5 pyramids (Fig. 1A,B). In layer 6 and in the lower portion of layer 5 (5b) fluorescence was absent.

**Figure 1.**
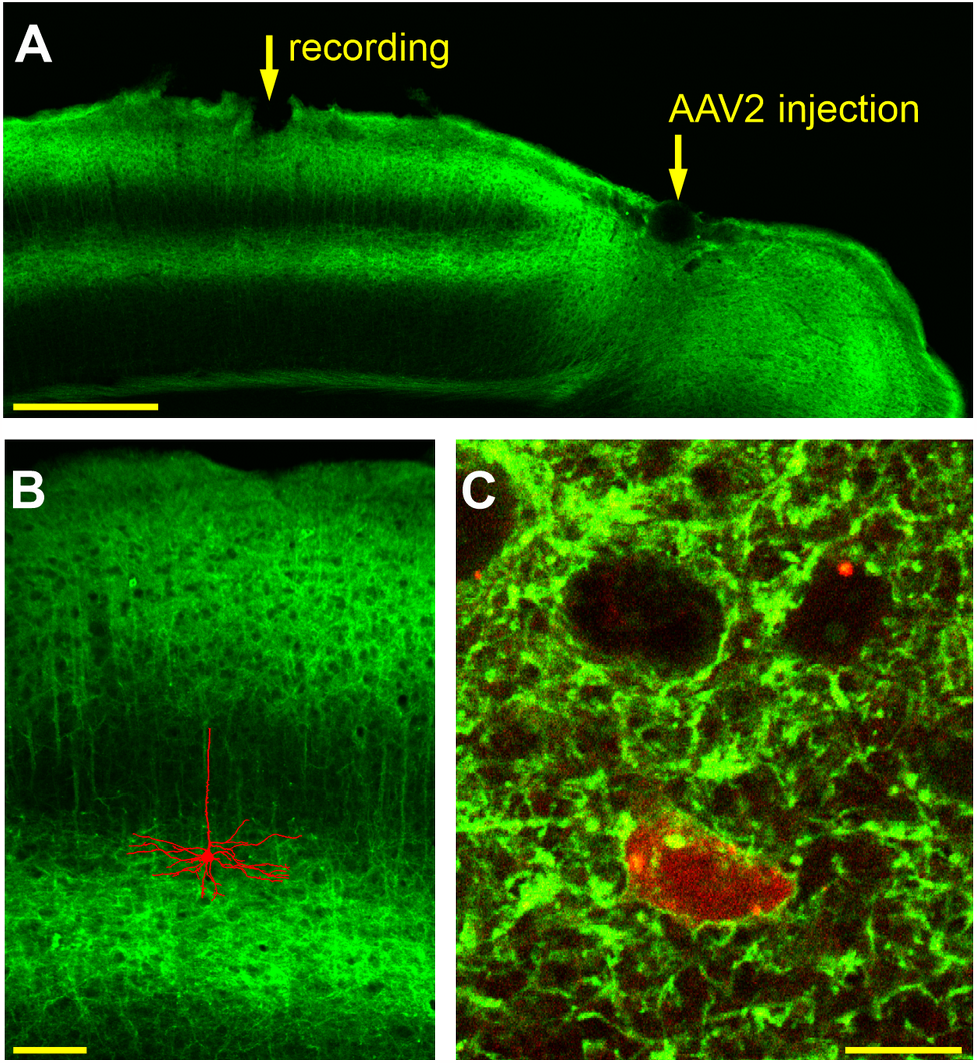
Expression of oChIEF-Venus in mouse visual cortex. A – Frontal section of the mouse visual cortex after transduction with the AAV2-CamKII-oChIEF-Venus virus, with the sites of virus injection and electrophysiological recording (2 weeks after the injection) indicated by the arrows. Note strong fluorescence in all cortical layers around the recording site, but with increasing distance from the injection site only in layers 1, 2/3, the upper portion of layer 5 and in the white matter. B – Pyramidal morphology of a neuron stained with neurobiotin during recording and then visualized with Streptavidin-Rhodamin. The image was superimposed with an image of a neighboring slice from the same animal showing the oChIEF expression pattern. Both images were taken using a confocal microscope, automatically traced and scaled. C – Soma of a neuron stained during recording with neurobiotin and visualized with Streptavidin-Rhodamine. A single confocal section. Green fluorescence of the somatic membrane is indicative of oChIEF-Venus expression. Calibration bars: A – 500 µm, B – 100 µm, C – 10 µm.

In 4 experiments neurons expressing oChIEF, as verified by the ability of light pulses to evoke action potentials, were labelled for morphological identification. In such experiments neurobiotin was added to the electrode solution and electroporated into the cell at the end of the recording. Out of 4 stained cells, 3 demonstrated typical morphology of layer 5 pyramidal neurons (Figure 2B), and one was a layer 2 neuron. Morphological analysis carried out using confocal microscopy showed that all cells stained during the recording with neurobiotin and visualized with Rhodamine, showed green fluorescence in the near-membrane region (Fig. 2C). Thus, expression of the oChIEF-Venus construct in these cells is verified using both physiological and morphological approaches.

**Figure 2.**
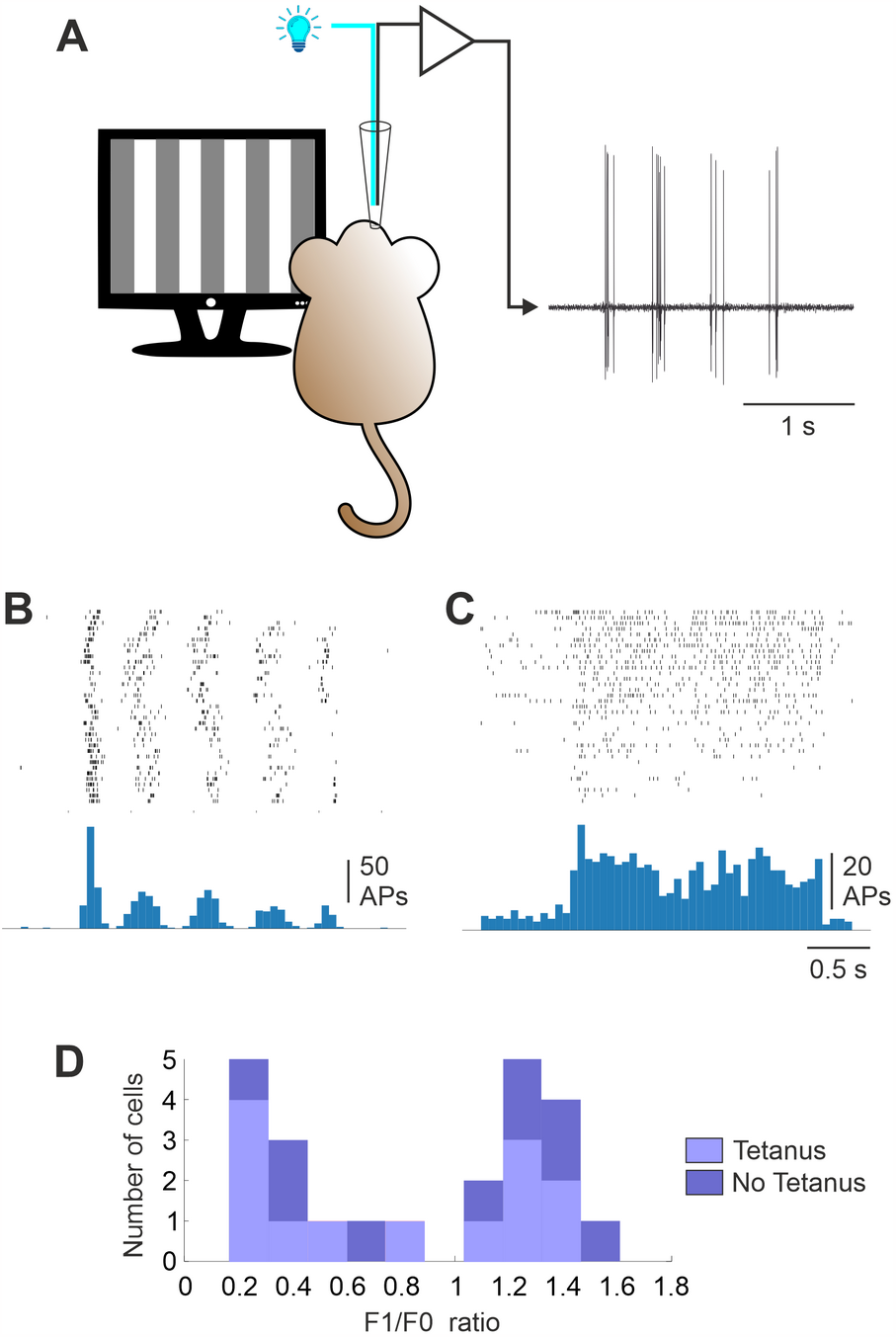
Responses of neurons with simple and complex receptive fields to visual stimulation. A – A scheme of the experiment. For local optogenetic stimulation an optical fiber was inserted into the recording glass electrode alongside with a silver wire for recording of neuronal activity. B, C – Examples of responses of a simple (B) and a complex (C) cell to a moving grating of optimal orientation. Raster diagrams of spike responses to 35 presentations of the moving grating and peri-stimulus time histograms plotted using the spikes from the rosters. Note the pronounced modulation of the response at the stimulus frequency (2 Hz) in a simple cell (B), but little modulation in a complex cell (C). D – Distribution of the ratio of the modulation component of the response to the average response amplitude (F1/F0 ratio) for cells in which optogenetic tetanization was applied (Tetanus group), and cells in the control (No Tetanus) group. Clearly bimodal distribution allows for a clear formal classification of cells as simple (F1/F0 > 1) or complex (F1/F0 < 1). Note an approximately equal number of simple and complex cells in the tetanus and no tetanus groups.

During the search for oChIEF expressing neurons, light pulses with a wavelength of 470 nm were continuously delivered through the optical fiber inserted into the recording electrode, while it was slowly lowered into the brain tissue. As soon as light-induced action potentials appeared (Fig. 2A) the electrode advancement was stopped, and visual stimulation started. Visual stimuli were whole-screen sinusoidal gratings (0.04 cpd, 2 Hz) moving in 12 different directions in a randomized order. Only neurons with clear responses to visual stimulation (see examples in Fig. 2B,C) were used for the experiments. We observed two types of responses to moving sine-wave gratings. Responses of the first type were characterized by a strong modulation of the visually-induced activity at the temporal frequency of the stimulus (Fig. 2B). Responses of the second type had a predominantly tonic increase of the spiking frequency during the stimulus presentation, but little modulation at the stimulation frequency (Fig. 2C). These distinct patterns of responses are characteristic features of visual cortex neurons with simple and complex receptive fields, well-described in different mammalian species and used for classification of cortical receptive fields [35–38]. For the formal classification of neurons, we calculated F1/F0 ratio using response to the optimal orientation. The envelope of the peri-stimulus time histogram was fitted by a sinusoid with the frequency of the stimulus. The F1/F0 ratio was calculated as the ratio of the amplitude of the fitted sine wave to the average amplitude of the response. The F1/F0 ratio of the recorded neurons showed a bimodal distribution, allowing a clear classification of neurons into simple (F1/F0 > 1, n=11) and complex (F1/F0 < 1, n=12) cells (Fig. 2D). The proportion of simple and complex cells in the experimental and control groups (see below) did not differ significantly (7 simple and 6 complex cells in the experimental group and 4 simple and 6 complex cells in the control group; Chi-square = 0.4343, p= 0.51).

Orientation tuning of a neuron was calculated using averaged responses (number of spikes) to stimuli of different orientations. To facilitate comparison across cells, orientation tuning of each neuron was normalized by the averaged response to all orientations of the moving gratings. Examples of typical orientation tunings of two neurons, plotted in polar coordinates, are shown in Figure 3A.

**Figure 3.**
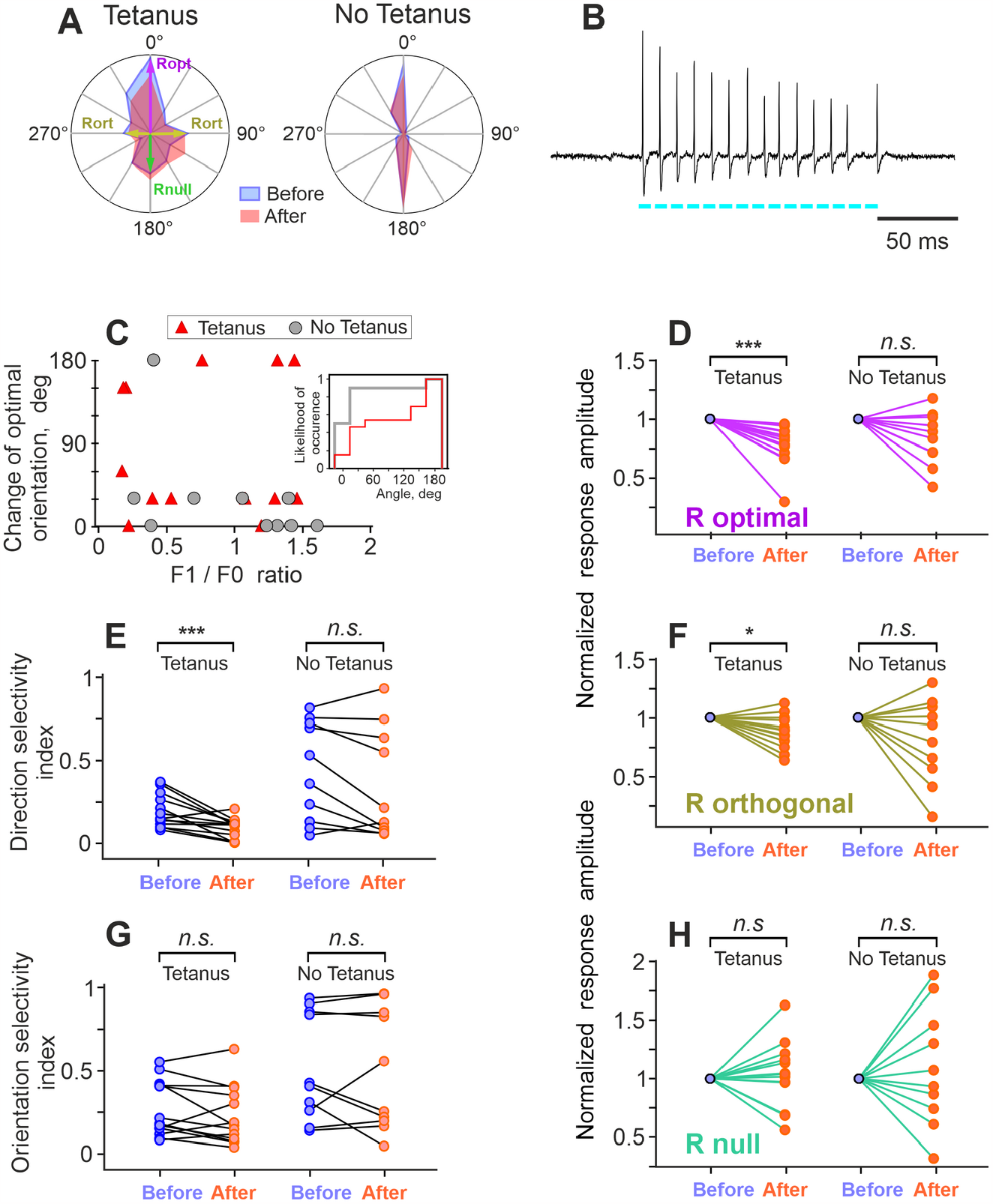
Optogenetic tetanization induces changes of responses to visual stimulation and functional properties of neurons in visual cortex. A – Examples of orientation tuning of a neuron subject to optogenetic tetanization (Tetanus group) and a neuron from control (No Tetanus) group. For both neurons, tunings before (blue) and after (pink) the tetanization, or a respective time interval without stimulation for the cell from control group, are shown. For the neuron in the Tetanus group, optimal (Ropt), orthogonal (Rort) and null orientations (Rnull) are indicated. B – A burst of action potentials induced by optogenetic tetanization in a neuron expressing oChIEF. The cyan bars below the trace show timing and duration of light stimuli delivered through the optical fiber inserted in the glass electrode. C – Changes of the optimal orientation after tetanization (or stimulation pause) plotted against the F1/F0 ratio. Note that changes of the optimal orientation occurred much more frequently after the tetanization (Tetanus group) than in the control (No Tetanus) group. Note also that the change in the optimal orientation did not depend on the F1/F0 ratio. Inset: Cumulative distributions of the optimal orientation change occurrence in the Tetanus (red) and No Tetanus (gray) groups. D, F, H – Changes of the normalized amplitude of responses to the stimuli of optimal (D), orthogonal (F), and null (H) orientation after tetanization (Tetanus, left parts of the plots) and in the control (No Tetanus, right parts of the plots) group. Response amplitude before tetanization (or before a pause in stimulation) was taken as 1; each pair of points connected by a line represents data from one cell. Color code as in A. E, G – changes in the indices of direction selectivity (E) and orientation selectivity (G) of visual cortex neurons after optogenetic tetanization (Tetanus) and in control group (No Tetanus). For those cells, in which the preferred orientation changed after the tetanization (or a pause in stimulation), the new, re-defined ‘optimal’ direction was used for calculation of parameters of ‘After’ responses.

After recording responses to moving gratings for 15-40 min, visual stimulation was stopped, and either an optogenetic tetanization of the neuron was performed (experimental group), or no action was taken (control group). Optogenetic tetanization consisted of 5 series of 10 trains of 8-10 pulses (8 ms, 50-100 Hz) of 470 nm light applied through the optical fiber inserted into the recording electrode. Each train of the optical tetanization evoked in the recorded neurons bursts of action potentials with a frequency of 50-100 Hz (Fig. 3B). Optogenetic tetanization lasted 5-6 minutes. After the optogenetic tetanization, or after 5-6 minutes without any stimulation in the control group, visual stimulation with moving gratings of different orientations was resumed and continued for at least one hour.

In the analysis, we first asked whether the optimal stimulus (grating of the optimal orientation moving in the preferred direction) changes after tetanization. Figure 3C shows changes of the optimal orientation of the moving grating after tetanization and in control group, plotted against the F1/F0 ratio. Note that because 30 deg was the step of orientations during tested, changes by 30 deg could have been spurious since ‘true’ optimal orientation could be between the tested ones. With this consideration in mind, the most frequent change was that optimal (strongest) response after tetanization was evoked by stimuli of the same orientation but moving in the opposite (previously null) direction (Fig. 3C).

In the tetanus group, the change in the optimal orientation of the stimulus occurred more frequently than in the control (no tetanus) group. There was significant difference between distributions of the optimal orientation changes in tetanus and no tetanus groups (p< 0.05, Anderson-Darling test; Fig. 3C inset). Respectively, averaged values of the change differed dramatically between the tetanus group (80.8±73.9 deg) and control group (30±54.8 deg). The change in the orientation of the optimal stimulus after tetanization did not correlate with the F1/F0 index (Fig. 3C). As described above, we classified cells with F1/F0 > 1 as simple, and with F1/F0 < 1 as complex (Fig. 2D). In the tetanization group, F1/F0 in simple cells was on average 1.3±0.14 (n=6), while in complex cells it was 0.35±0.22 (n=7).Next, we compared responses to each orientation of the grating before and after the tetanization (or after 5-6 minutes without stimulation in the control group). Optogenetic tetanization had differential effects on responses to different orientations.

After tetanization, the amplitudes of responses to the gratings of optimal orientation significantly decreased to 77.7±17.3% compared to the values before tetanization (p<0.001, n=13, Wilcoxon matched-pairs signed rank test; Fig. 3D, left panel). Responses to non-optimal (orthogonal to optimal) orientation decreased to 88.8±15% (p<0.0327, Fig. 3F, left panel). In the control (no tetanus) group, responses to neither the optimal orientation, nor to the non-optimal orientation changed significantly (83.5±25%, n.s., and 89.1±27.8%, n.s.; n=10; Fig. 3 D,F, right panels). Responses to the null direction of movement (grating of optimal orientation but moving in the opposite to preferred direction) did not change significantly in the two groups (113±35% for the experimental group, 111±47.6% for the control group, Fig. 3H). As a consequence of the decrease of responses to the optimal stimulus, but unchanged response to the stimulus moving in the null direction, there was a significant decrease in the directional selectivity index after optogenetic tetanization, which decreased to 51.5±38.2% of the values before tetanization, from 0.22±0.15 to 0.096±0.06, (p<0.0017, Wilcoxon matched-pairs signed rank test). In the control no tetanus group the direction selectivity index did not change (Fig. 3E). The orientation selectivity index did not change after tetanization, indicating that the amplitudes of responses to the optimal and to the orthogonal orientations of the stimuli decreased proportionally (Fig. 3G).

For those parameters that changed significantly after tetanization, we analyzed separately the changes among simple and complex cells. There was no significant difference between simple (n=6) and complex (n=7) cells in the decrease of the responses to neither the optimal orientation (75.4±10% and 79.6±3%), nor the nonoptimal (orthogonal to the optimal) orientation (82.7±12% and 93.7±15%), nor the direction selectivity index (44.5±50% and 57.4±27%) (n.s., Student’s t-test in all three comparisons). This result confirms the validity of pooling together the data for simple and complex cells.

In the next step of the analysis, we compared the tuning of cells to orientation and direction of stimuli before and after tetanization. For calculation of averaged orientation tuning, we recentered the tuning of each cell so that the orientation of the optimal stimulus was at 0 deg, and then mirrored the tuning plotted in polar coordinates along the optimal orientation. Then we averaged the responses to stimuli ±30 deg, to stimuli ±60 deg, and so on. Figure 4 A,B shows the resulting “half” tuning curves plotted in Cartesian coordinates for the tetanus and no tetanus groups. As described above, after tetanization there was a significant decrease in the amplitude of responses to the optimal and orthogonal orientations (Fig. 4A). In addition, there was an increase in the amplitude of responses to stimuli oriented 120 and 150 degrees from the optimal (0°: 78.1 ±27% p = 0.0098, 90°: 87.9 ± 31% p = 0.0406, 120°: 125 ± 29% p = 0.0406, 150°: 116± 26% p = 0.0406, Wilcoxon rank test adjusted for multiple comparison using false discovery rate method). In the control (no tetanus) group, no significant changes of responses to stimuli of any orientation were found (Fig. 4B). To represent changes of the directional tuning, we redraw the “whole” tuning curve from Figure 4A in polar coordinates (Fig. 4C). As a result of the above changes after optogenetic tetanization: decrease of the responses to stimuli of optimal and orthogonal orientation, no changes of responses to null orientation, and an increase of responses to 120 and 150 deg; the curve became more round-shaped, i.e., the cells became less selective.

**Figure 4.**
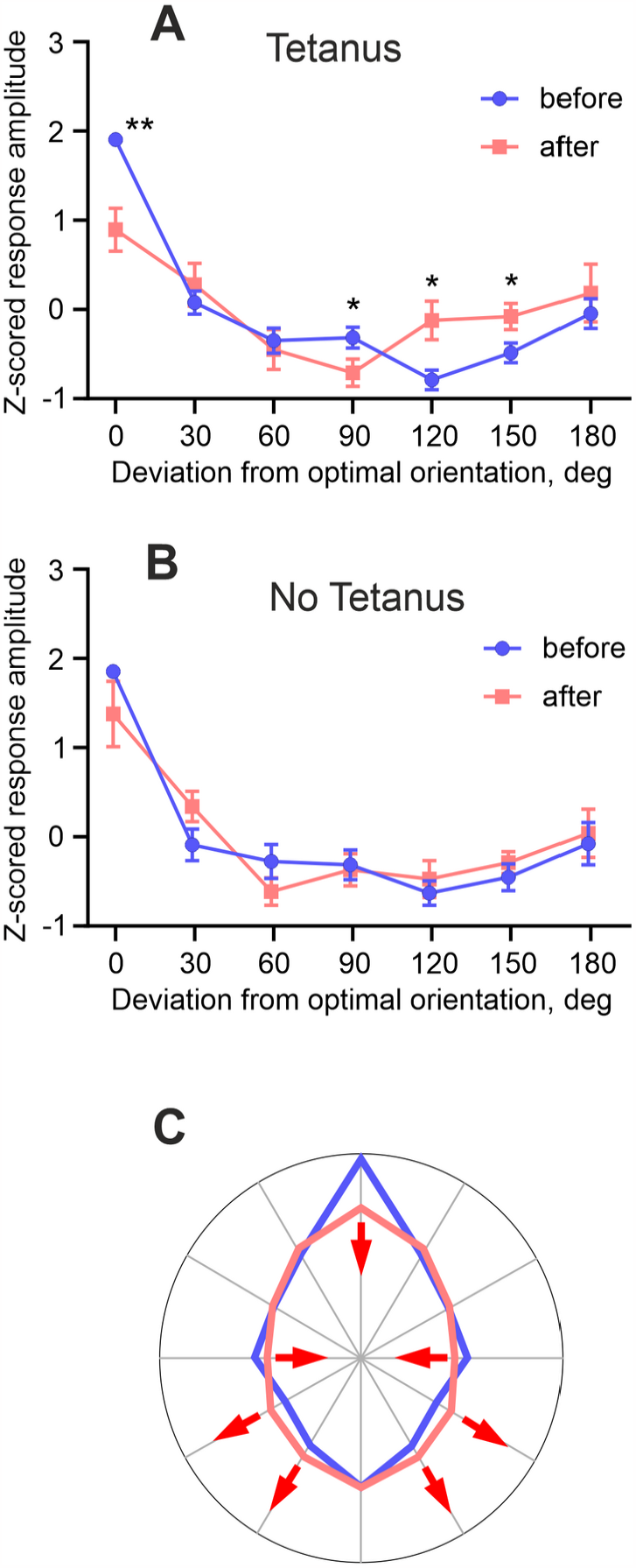
Optogenetic tetanization induces changes of the tuning of neurons to orientation and direction of movement of visual stimuli. A, B – Averaged tuning of responses of cortical neurons to orientation before (blue) and after (pink) tetanization (A) or before and after a stimulation pause in control group (B). Each point represents average responses to the grating of one orientation moving in two opposite directions. To facilitate comparison, all values were Z-scored. * p<0.05; ** p < 0.01. C – A schematic representation of changes of responses to different directions of movement of visual stimuli, induced by optogenetic tetanization of mouse visual cortex neurons.

## DISCUSSION

Results of our study show that high-frequency bursts of action potentials induced in visual cortex neurons using direct optogenetic stimulation (optogenetic tetanization) can cause long-term changes of neurons’ responses to visual stimuli. Changes of responses to visual stimuli reflect changes of synaptic inputs to the neuron, and because optogenetic tetanization is a purely postsynaptic protocol applied without presynaptic activation, such changes were mediated by heterosynaptic mechanisms. These results provide clear evidence for heterosynaptic plasticity in the whole brain *in vivo*. Moreover, optogenetic tetanization had distinct effects on responses to stimuli of different orientations, indicating that heterosynaptic plasticity affected neuronal inputs non-uniformly.

### Homosynaptic and heterosynaptic plasticity, terminology

The terms homosynaptic and heterosynaptic are used in somewhat different ways in the field of plasticity studies in mammalian preparations and in invertebrate research. Here we use this terminology in a way that is conventional for plasticity research in mammals. *<u>Homosynaptic</u>* is plasticity induced at synapses which were presynaptically activated during the induction. It can be induced by protocols such as diverse patterns of afferent tetanization, and pairing protocols: stimulation of weak synaptic inputs together with activation of strong inputs or postsynaptic depolarization [2,39,40]. If activity at both presynapse and postsynapse is required for the induction, homosynaptc plasticity is associative, classical example here is NMDA-dependent plasticity [40]. If plasticity induction depends only on strong presynaptic activity but does not require activity (depolarization and spiking) of the postsynaptic cell, homosynaptic plasticity is non-associative, such as at mossy fiber-CA3 synapses in the hippocampus [41]. *<u>Heterosynaptic</u>* is plasticity that does not require activity in the presynapse during the induction. In the field of synaptic plasticity in the hippocampus, this term was introduced by Lynch and colleagues [5], who found that tetanization-induced LTP at synapses terminating at apical dendrites of CA1 pyramids is accompanied by LTD at synapses on the basal dendrites, and vice versa, tetanus-induced LTP at the basal dendrites was accompanied by LTD at the apical dendrites. Because synapses undergoing LTD were not stimulated during the induction, this form of LTD was called heterosynaptic. Moreover, heterosynaptic plasticity, both LTP and LTD, could be induced in cortical neurons by episodes of strong postsynaptic activity alone, without stimulation of any synapses during the induction [15,16,43,22–28,42]. Local forms of heterosynaptic plasticity include induction of LTP and LTD around the location of the tetanized synapses, forming a Mexican hat-shaped profile of plastic changes [17–19], and spread of plastic changes to non-stimulated synapses of the same neuron, or even to closely located but non-stimulated neurons [20,21]. A distinctive feature of these forms of plasticity is that they are induced at synapses which were not active during the application of the induction protocol, and thus are heterosynaptic (see [4,44] for review).

In invertebrate research the terms homosynaptic and heterosynaptic are used in a different way. Kandel and Tauc [6,7], studying facilitation of synaptic transmission in *Aplysia*, introduced the term heterosynaptic to describe pairing of weak stimulation of one pathway with a strong stimulation of another pathway (‘heterosynaptic pairing’). Such pairing may lead to a short-living (9 minutes on average, up to 40 minutes in extreme cases) facilitation of responses in the weakly stimulated pathway. Further studies of gill-withdrawal in Aplysia demonstrated the importance of serotonergic modulation during the strong stimulation for behavioral reaction and heterosynaptic facilitation [45]. Later, these ideas were generalized, and modulatory input-dependent plasticity was referred to as heterosynaptic, while Hebbian activity-dependent as homosynaptic [8]. The term homosynaptic plasticity was also used [8] as a synonym of the property of cooperativity of LTP in mammalian plasticity research [40]. Notably, in the original terminology [6,7], heterosynaptic pairing in *Aplysia* research roughly corresponds to pairing induced homosynaptic plasticity, and also to the features of cooperativity and associativity of LTP in the hippocampus [40].

### Plasticity of responses and receptive fields in visual cortex neurons

Prior research on plasticity of visual responses and receptive fields of neurons in visual cortex *in vivo* was focused on modifications induced by associative pairing procedures. As we demonstrated in our prior work, pairing visual stimulus of non-optimal orientation with optogenetically induced firing of pyramidal neurons in mouse visual cortex induced long-term changes of the response properties of the stimulated cell: responses to previously non-optimal (paired) stimulus increased while responses to the originally preferred orientation of the stimulus decreased [11]. In cat visual cortex, pairing of flashing gratings of near-optimal and optimal orientation led to a significant shift of orientation preference [46], and pairing of stimulation of non-responsive locations neighboring to the neuron’s receptive field with stimulation of receptive field center could lead to increase of the RF toward the stimulated site [9]. Pairing visual stimulation with intracellularly evoked spikes in neurons of rat visual cortex induced bidirectional modifications of responses. Importantly, responses evoked by stimulation of unpaired locations could change too, indicative of heterosynaptic changes [10]. Heterosynaptic changes were also reported in a study using imaging of calcium signals in spines of mouse visual cortex neurons [31]. Pairing visual stimulation that activated a spine with postsynaptic spiking induced LTP in the activated spine, and heterosynaptic LTD at some of the other spines on the same neuron. With a reservation that absence of presynaptic activation at inputs which were not intentionally stimulated is difficult to control with strong visual stimulation and large subthreshold receptive fields, these results indicate that heterosynaptic plasticity can be induced in visual cortex neurons of rat and mouse.

### Paradigm of optogenetic tetanization to study heterosyanptic plasticity

Advantage of our approach to study heterosynaptic plasticity using optogenetic tetanization is that, in the absence of visual stimulation during the induction, the probability of repeated presynaptic activation of neuron’s inputs leading to ‘spurious STDP’ pairing with postsynaptic spikes is negligibly small. Indeed, plasticity induction requires consistent and repetitive pre-before-post activation for LTP, or post-before-pre for LTD, for >30 times [3,10,47]. With typical spontaneous firing rates <5 Hz, a presynaptic spike could hit a ∼50-100 ms STDP window [2,10,48] of light-induced postsynaptic spikes, however the probability that such coincidence will occur >30 times consistently only in LTP (or only in LTD) window is extremely low. Rather, presynaptic spikes that occasionally hit the LTP and LTD windows will do so at random, cancelling the effects of each other. Therefore, we consider such scenario of ‘spurious STDP’ unrealistic, and attribute changes of responses after optogenetic tetanization to heterosynaptic plasticity.

The paradigm of optogenetic tetanization, implemented in our study, is a transfer to *in vivo* conditions of the protocol of intracellular tetanization, which is established in brain slice preparations and leads to bi-directional long-term plastic changes of synaptic inputs [16,23,24] (see [4,44] for review). While the high-frequency bursting of cells induced by pulses of depolarization current or light is an artificial pattern, it has clear correlates to some patterns of activity in cortical neurons observed *in vivo*. Pyramidal neurons in rat hippocampus can generate bursts of action potentials with frequencies of up to 200 Hz during spontaneous activity [49]. In cat visual cortex, patterns of high frequency activity comparable to optogenetic tetanization can be produced by optimal visual stimulation [37]. Similar patterns of spiking are also typical for slow-wave sleep, with periodic transitions of cortical neurons between active (up) and silent (down) states [50–52]. Absence of sensory input from the thalamus and predominantly intracortical origin of activity during slow-wave sleep [53] adds to this similarity. Thus, the pattern of spiking induced by optogenetic tetanization is similar to some naturally occurring patterns of activity in cortical neurons, especially to those during slow-wave sleep, which might therefore promote induction of heterosynaptic plasticity.

### Possible mechanisms and functional consequences of plasticity induced by optogenetic tetanization

Our present results show that after optogenetic tetanization the most pronounced changes were in the directional selectivity of neurons, and often preferred direction of movement was changed to the opposite. This was due to a decrease of responses to stimuli of optimal orientation moving in preferred direction, but no changes of responses to the opposite (null) direction of movement. Because directional selectivity is created by the interaction of excitatory and inhibitory inputs [54–56], such changes could be mediated by depression of excitatory inputs or potentiation of inhibitory inputs, or a combination of both. Indeed, in slices heterosynaptic plasticity can be induced at both excitatory synapses [14,16,23,25] and at inhibitory GABA-ergic synapses to layer 5 neurons [43]. Present results do not allow us to differentiate between these possibilities. Disentangling the contribution of a decrease of excitation and/or an increase of inhibition requires further studies, e.g. employing intracellular recordings.

A decrease of responses to optimal and orthogonal orientations, no change of responses to null direction and an increase of responses to oblique orientations after optogenetic tetanization made the tuning of cells more round-shaped, and responses less selective (scheme in Figure 4C). Several aspects of these results are worth noting. First, the decrease of strong (optimal) responses and increase of relatively weak response to oblique orientations has a parallel with weight-dependence of heterosynaptic plasticity, observed *in vivo* [10] and extensively studied in slices and computational models [4,14,24,25,27]. Second, the weight-dependence of response changes induced by heterosynaptic plasticity might help to prevent excessive increase of responses to optimal stimuli and runaway decrease of responses to other than optimal stimuli due to runaway potentiation or depression of respective inputs due to plasticity governed by Hebbian-type rules. Third, theoretical studies using information theory approach show that there is an optimal width of tuning of individual neurons for encoding of three or more features of visual stimuli in population activity [57,58]. Such optimal width allows each neuron of a population to encode the maximal amount of information about the visual stimuli. With the tuning width outside the optimal range, the amount of information a neuron can encode is decreasing. Since neurons in primary visual cortex encode multiple features of visual stimuli, such as orientation, direction and speed of motion, size, etc, we hypothesize that response changes induced by heterosynaptic plasticity may help to keep the width of orientation and direction tunings around such optimal values. Thus, we suggest that heterosynaptic plasticity could play a role not only in stabilizing the orientational and directional tunings of individual neurons, but also in maintaining their width within the range optimized for high efficiency of encoding of properties of visual stimuli in population activity.

To summarize, we show that, in neurons of mouse visual cortex, optogenetic tetanization can induce changes of responses to visual stimuli. This provides evidence for heterosynaptic plasticity in the whole brain *in vivo*. Based on the observed response changes we hypothesize that homeostatic role of heterosynaptic plasticity, in addition to its importance in preventing runaway dynamics of synaptic weights and synaptic drive of individual neurons, extends to the level of neuronal population encoding, by stabilizing the tuning of neuronal responses to features of visual stimuli and keeping the tuning width in the range that is optimal for encoding of multiple features. Finally, because of the similarities between the patterns of activity during slow-wave sleep and optogentic tetanization, we suggest that heterosynaptic plasticity may act as one of the mechanisms mediating homeostatic function of the slow wave sleep.

